# Acquired deficiency of the peroxisomal enzyme enoyl-CoA hydratase/3-hydroxyacyl CoA dehydrogenase is a metabolic vulnerability in hepatoblastoma

**DOI:** 10.1101/2020.08.24.265421

**Authors:** Huabo Wang, Xiaoguang Chen, Marie Schwalbe, Joanna E. Gorka, Jordan A. Mandel, Jinglin Wang, Jie Lu, Eric S. Goetzman, Steven Dobrowolski, Edward V. Prochownik

## Abstract

Metabolic reprogramming provides transformed cells with proliferative and/or survival advantages. However, capitalizing on this therapeutically has been only moderately successful due to the relatively small magnitude of these differences and because cancers may re-program their metabolism to evade metabolic pathway inhibition. Mice lacking the peroxisomal bi-functional enzyme enoyl-CoA hydratase/3-hydroxyacyl CoA dehydrogenase (Ehhadh) and supplemented with the 12-carbon fatty acid lauric acid (C12) accumulate dodecanedioic acid (DDDA), a toxic C12 metabolite that causes hepatocyte necrosis and acute liver failure. In a murine model of pediatric hepatoblastoma (HB), down-regulation of Ehhadh also occurs in combination with a more general suppression of mitochondrial β- and peroxisomal ω-fatty acid oxidation (FAO) pathways. HB-bearing mice provided with C12 and/or DDDA-supplemented diets survived significantly longer than those on standard diets. The tumors also developed massive necrosis in response to short-term DDDA supplementation. Reduced Ehhadh was noted in murine hepatocellular carcinomas (HCCs) and in substantial subsets of human cancers, including HCCs. Acquired DDDA resistance was not associated with Ehhadh re-expression but was associated with 129 transcript differences ~90% of which were down-regulated in DDDA-resistant tumors and ~two-thirds of which correlated with survival in several human cancers. These transcripts often encoded components of the extracellular matrix suggesting that DDDA resistance arises from its reduced intracellular transport. Our results demonstrate the feasibility of a metabolic intervention that is non-toxic, inexpensive and likely compatible with traditional therapies. C12 and/or DDDA-containing diets could potentially be used to supplement other treatments or as alternative therapeutic choices.

## Introduction

Cancer cells undergo numerous metabolic adjustments to acquire and maintain proliferative and survival advantages and these represent essential features of the transformed state (1–7). The classic example of metabolic reprogramming is the Warburg effect, whereby the anaerobic process of glycolysis continues or is even accelerated under aerobic conditions (7–9). The excess of glycolytic intermediates provided by Warburg-type respiration is believed to serve as a ready source of the reducing equivalents, ribose sugars, nucleotides and amino acids needed to sustain tumor growth. ATP production is also increased during Warburg-type respiration and may even exceed the levels provided by oxidative phosphorylation (Oxphos), the down-regulation of which is a prominent feature of many cancers (8,10,11). Yet despite its diminished prominence, mitochondrial metabolism may continue to be an important source of TCA cycle-derived anabolic precursors such as acetyl coenzyme A (AcCoA), α-ketoglutarate and succinate (1,12,13). Indeed the up-regulation of pathways that provide anaplerotic sources of these TCA cycle substrates is another important metabolic feature of many cancers (1,2,5,14,15). Many studies have attempted, with variable success, to leverage some of the more prominent metabolic differences between normal and transformed cells for therapeutic benefit (1,4,9,14,16,17).

We previously observed that murine models of pediatric and adult liver cancer, namely hepatoblastoma (HB) and hepatocellular carcinoma (HCC), as well as their human counterparts, also remodel their metabolism and become more reliant on glycolysis than on fatty acid oxidation (FAO). Consistent with this, total mitochondrial mass and transcripts encoding virtually all enzymes in the mitochondrial-based β-FAO pathway are markedly down-regulated (18–22).

In addition to β-FAO, most cells also engage in non-energy-generating ω-oxidation, one purpose of which is to catabolize medium chain fatty acids that are too small to be transported by the fatty acid carrier carnitine palmitoyltransferase 1A (Cpt1a) and too large to enter mitochondria passively. ω-oxidation initiates in the endoplasmic reticulum via reactions involving the cytochrome P450 members Cyp4a10 (CYP4a11 in humans) and Cyp4a14, with subsequent catabolism occurring via peroxisomal β-oxidation. Some of the end products of this pathway such as succinate and AcCoA, can be utilized for anabolic purposes whereas others, such as adipate and suberate, are excreted.

12-carbon long lauric acid (hereafter C12) is the main fatty acid component of coconut oil and is metabolized primarily by the above ω-oxidation/peroxisomal pathway (23). Among its intermediate products is dodecanedioic acid (DDDA), a highly toxic dicarboxylic acid that is further metabolized by the bifunctional enzyme enoyl-CoA hydratase/3-hydroxyacyl CoA dehydrogenase (Ehhadh) (24). In contrast to normal mice, which readily tolerate either C12- or DDDA-enriched diets, *ehhadh-/-* mice accumulate high levels of DDDA, develop hepatic inflammation and necrosis and succumb within days to acute liver failure (25,26).

In addition to down-regulating mitochondrial β-oxidation, we show here that experimental murine HBs and HCCs also markedly suppress the ω-oxidation/peroxisomal pathway (18,20,21). This suggested that these tumors and perhaps other cancers, possess a metabolic vulnerability in the form of C12 and/or DDDA intolerance analogous to that of *ehhadh-/-* mice. Indeed, we also show that the lifespans of HB-bearing mice can be significantly extended by dietary supplementation with C12 and/or DDDA provided shortly after tumor initiation. Similarly, large pre-existing tumors develop massive necrosis in response to short-term implementation of DDDA-enriched diets. Several different human cancer types, including HB and HCC, also show striking down-regulation of *ehhadh* expression suggesting that these too might be sensitive to C12 and/or DDDA. Collectively, our studies identify a novel metabolic susceptibility that could be targeted via relatively simple means and with minimal toxicity without compromising more traditional chemotherapeutic options.

## Results

### C12- and/or DDDA-enriched diets extend the lifespan of HB-bearing mice

HBs were generated by hydrodynamic tail vein injection (HDTVI) of Sleeping Beauty (SB) vectors encoding a 90 bp in-frame oncogenic deletion mutant of β-catenin [hereafter Δ(90)] and a missense mutant (S127A) of yes-associated protein (hereafter YAP^S127A^), the terminal effector of the Hippo signaling pathway (19–22,27). We refer to these HBs hereafter as Δ(90) tumors. After allowing four weeks for tumor initiation, the animals were divided into four dietary cohorts: 1. standard diet, 2. standard diet+10% (w/w) C12 in the form of glycerol trilaurate (C12 diet), 3. standard diet+10% (w/w) DDDA (DDDA diet) and, 4. standard diet+C12 and DDDA (10% w/w each) (C12+DDDA diet). As previously reported, median survival in the former group was 14.2 wk (19–22,27) (Fig. 1A). In contrast survival among each of the other groups was significantly longer (P=1.4×10^−8^) although they were similar among themselves (range 22.8-28.6 wks, P=0.56-0.99). These results show that two structurally distinct substrates of the ω-oxidation/peroxisomal pathway each significantly extended longevity in this model of aggressive HB (28–30).

**Figure 1.**
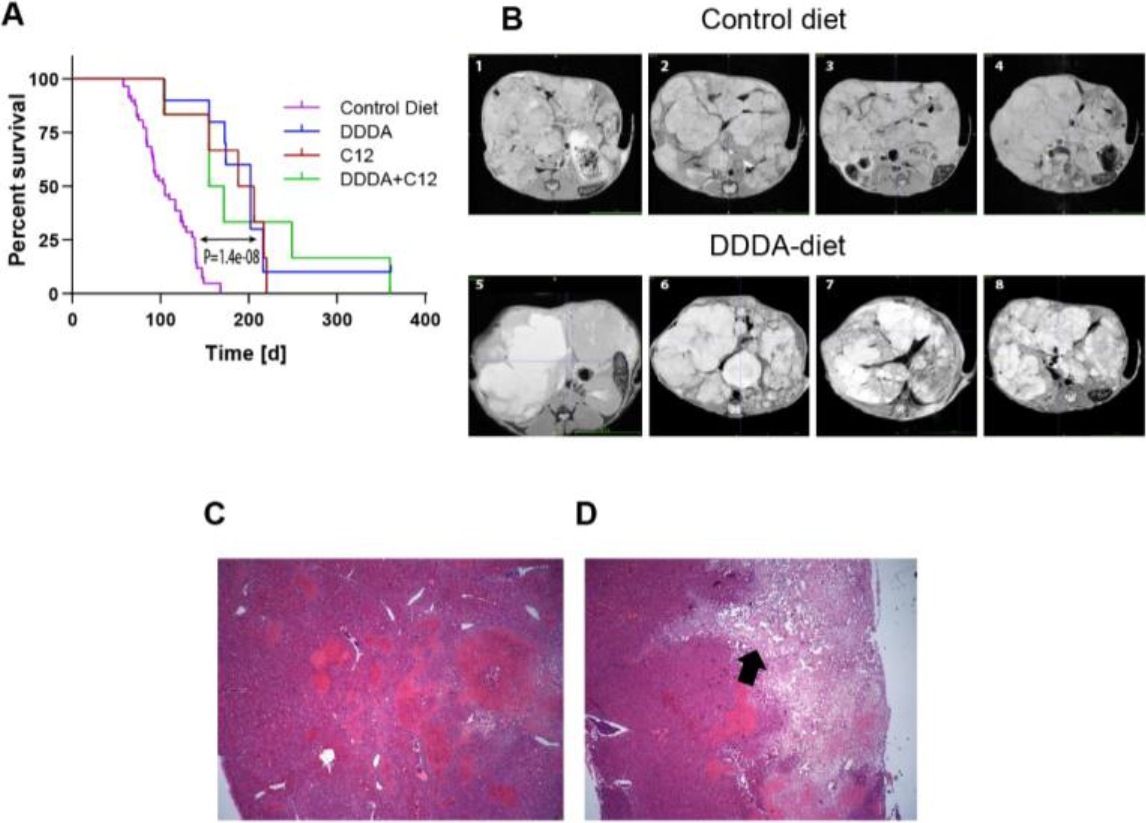
C12 and DDDA diets impair long-term HB growth and induce massive necrosis. *A.* Survival curves of HB-bearing mice. Each of the indicated cohorts was injected with SB vectors encoding Δ(90) and YAP^S127A^ (19–22,27). One week later the indicated diets were initiated for the remainder of the study. Kaplan-Meier survivals for each group were determined by the log-rank test. *B.* Tumors were generated as described in (A) except that the mice were then maintained on standard diets for eight wks. Half the mice were then switched to DDDA diets and the remaining mice were maintained on standard diets. MRIs were performed on each group at wk 11. Representative axial sections from four mice in each group are shown*. C.* Low-power magnification (5X) of a typical H&E stained section from a standard diet HB obtained at the time of MRI. *D.* Similar H&E section of a typical HB from a mouse maintained for three wks on a DDDA diet as described in *B.* A large area of necrosis extending from the tumor surface into the parenchyma is indicated by the arrow.

To determine how the above interventions impacted the growth of more advanced HBs, we generated additional Δ(90) tumors, initiated DDDA diets in half the animals after tumors had attained a moderately large size at 8 weeks and then evaluated the tumors 3 wks later by MRI. As expected, mice maintained on standard diets had large tumors that displaced adjacent organs. They also succumbed by week 12-13 with 10-14 gram tumors (Fig. 1B)(19,20,22). In contrast, tumors from mice maintained on DDDA diets showed areas of extensive necrosis that was confirmed by histologic examination of H&E-stained tumor sections (Fig. 1C and D). Control HBs contained densely packed regions of small blue cells with the crowded fetal histology previously reported for Δ(90) tumors, representing the most common HB subtype (19–22,27,29). While histologically similar, DDDA diet tumors contained large areas of consolidative necrosis, which penetrated deeply into the adjacent parenchyma. Together with the previous results, these findings indicate that C12 and/or DDDA slow HB progression and promote regression by inducing extensive tumor cell death.

### Tumors down-regulate mitochondrial- and peroxisomal FAO

Δ(90) HBs coordinately down-regulate mitochondrial mass by ~80% as they switch their primary mode of energy generation from mitochondrial β-FAO to glycolysis (19–22,30). We observed a similar down-regulation of transcripts related to endosomal ω- and peroxisomal FAO (hereafter ω-/peroxisomal FAO), which is largely responsible for the metabolism of fatty acids such as C12 (24,25,31) and found that, as a group, they were 3.1-fold down-regulated (P= 2.6×10^−12^, Fig. 2A and B). Among these transcripts were those encoding Ehhadh (7.7-fold down-regulated, P=1.11×10^−16^) and the microsomal P450 cytochromes Cyp4a10 and Cyp4a14, which catalyze the first step in the catabolism of C12 to various dicarboxylic acids (13.1-fold and 35.8-fold down-regulated, respectively, P=4.31×10-7, 6.98×10^−11^)(32). This paralleled an equally marked decline at the protein level (Fig. 2C).

**Figure 2.**
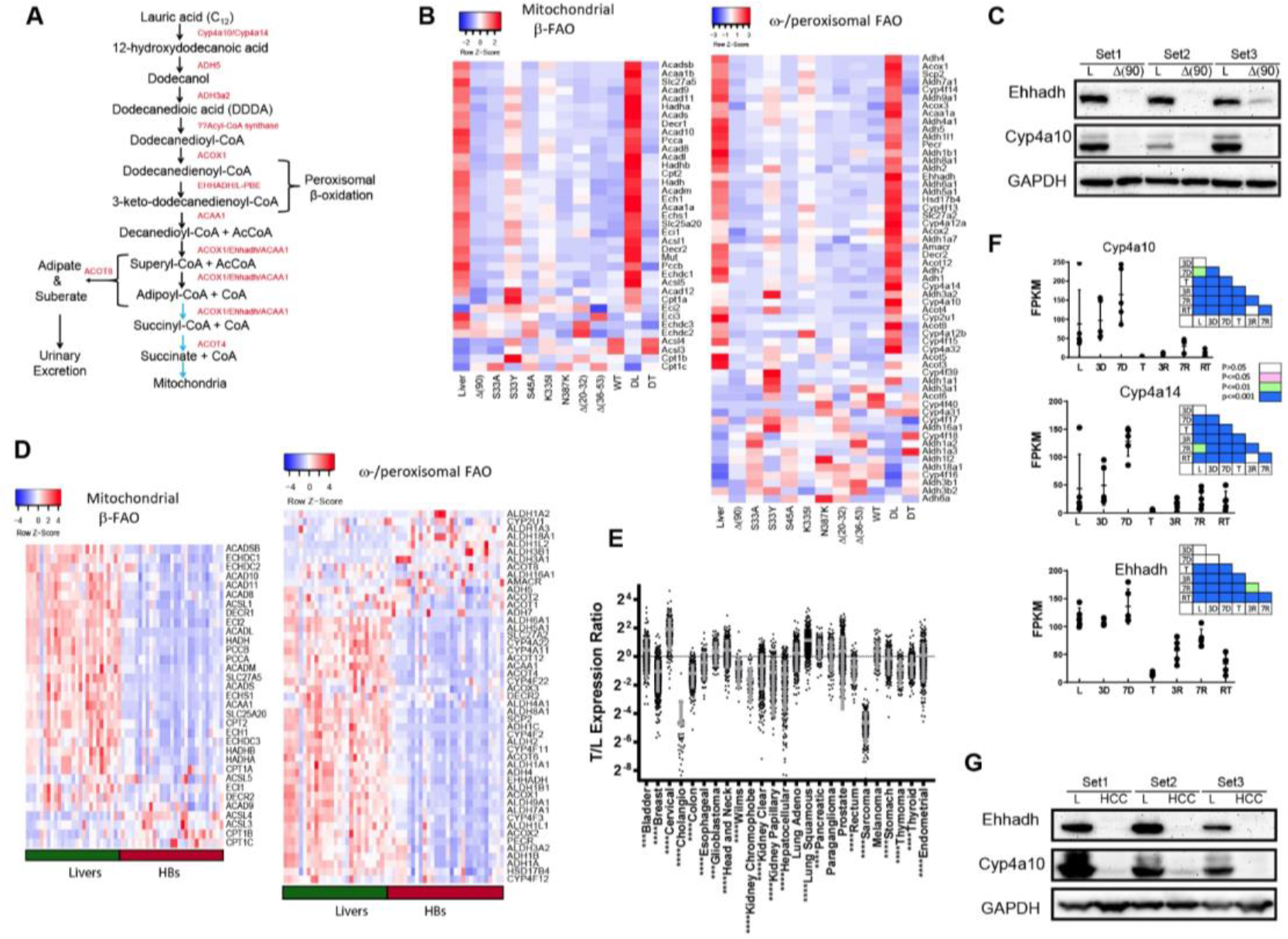
Parallel down-regulation of the β -FAO and ω -/peroxisomal-FAO pathways in murine and human cancers. *A.* The pathway of C12 metabolism. The initial step of ω -oxidation occurs in endosomes and is catalyzed by Cyp4a10 and Cyp4a14. The resulting dicarboxylic acids, including DDDA are further catabolized in this compartment or in the peroxisome. The ultimate products, adipate and suberate are excreted in urine whereas succinyl-CoA and succinate supply the TCA cycle or enter biosynthetic pathways. *B.* Heat maps showing transcript levels for mitochondrial β-FAO and ω-/peroxisomal-FAO pathways in livers and Δ(90) HBs maintained on standard and DDDA diets (left-most and right-most columns, respectively). DL and DT=livers and tumors from DDDA-diet cohorts. In addition, transcripts are shown from standard diet-maintained tumors generated by other previously described patient-derived β-catenin mutants (22)*. C.* Ehhadh and Cy4a10 protein expression from three independent sets of livers and Δ(90) β-catenin mutant-generated HBs, all from mice maintained on standard diets. *D.* Heat maps as described in *B* generated from 24 human HBs. *E*. Expression of Ehhadh transcripts from the indicated human tumors relative to that of matched control tissues from TCGA. *F.* Absolute expression (FPKM) of transcripts encoding Cy4a10, Cyp4a14 and Ehhadh in tissues from mice bearing a doxycycline-responsive human Myc transgene. Results are taken from RNAseq data published in Dolezal et al (18). L: control (Myc-suppressed) livers prior to Myc induction; 3d and 7d: livers without obvious tumors 3 days and 7 days after withdrawal of doxycycline and the induction of Myc; T: tumors samples approximately 4 wks after the induction of Myc; 3R and 7R: regressing tumors 3d and 7d following Myc silencing after doxycycline reinstatement. RT: recurrent tumor. Initial tumors were allowed to regress for approx. 3 months at which point no residual tumor could be detected (18,34). Recurrent tumors were then induced by the removal of doxycycline and sampled after 3-4 wks of tumor re-growth. *G.* Expression of Ehhadh and Cyp4a10 in three sets of control livers and Myc-driven initial HCCs from F.

In concert with YAP^S127A^, numerous other patient-derived β-catenin mutants are tumorigenic in mice (22). These grow at different rates and display diverse histologic subtypes, in some cases more closely resembling HCCs than HBs. However, like Δ(90) tumors, they all broadly down-regulated transcripts encoding β-FAO and ω-/peroxisomal FAO enzymes (Fig. 2B and (22). The suppression of the latter pathway is thus widespread, occurs in parallel with mitochondrial β-FAO-related transcript down-regulation and is independent of tumor growth rate. The transcriptional profiles of 24 previously reported primary HBs (33) also showed that, like the case for murine HBs, these tumors down-regulated both β-FAO and ω-/peroxisomal pathways (Fig. 2D).

Among the 33 tumor types in The Cancer Genome Atlas (TCGA) for which a sufficient number of samples plus matched control normal tissues were available, we identified at least six tumor types where ω-/peroxisomal-FAO transcripts as a group were significantly down-regulated, although considerable variability was seen among individual tumors (Supplementary Fig. 1). Those containing subsets with particularly low expression of Ehhadh included HCCs, cholangiocarcinomas and sarcomas (Fig. 2E).

Aggressive murine liver cancers resembling poorly differentiated HCCs arise in response to the conditional over-expression of c-Myc (Myc) (18,34). These tumors regress rapidly and completely following Myc silencing and recur following its re-expression. We followed Cyp4a10, Cyp4a14 and Ehhadh transcripts across this time course. In contrast to Ehhadh transcripts, which remained unchanged during the earliest stages of tumor induction and prior to their actual appearance, Cyp4a10 and Cyp4a14 transcripts were initially induced in a time-dependent manner (Fig. 2F). However, all three transcripts fell to near undetectable levels in the tumors that subsequently appeared, thus mimicking their behavior in HBs (Fig. 2B). A partial normalization of transcripts, particularly those encoding Ehhadh was seen during regression. Recurrent tumors, like the initial ones, also expressed low levels of all three transcripts. Finally, as had been seen with HBs, Ehhadh and Cyp4A10 protein levels were virtually undetectable in tumors (Fig. 2G). Taken together, these results show that the parallel down-regulation of the mitochondrial β-FAO and ω-/peroxisomal-FAO pathways occurs in both murine HBs and HCCs, that a similar down-regulation occurs in human HBs and that ω-/peroxisomal-FAO pathway suppression is a frequent albeit variable finding across some human cancer types, including HCCs.

β-FAO- and ω-/peroxisomal-FAO-related transcripts tended to correlate in a number of human cancers and in at least six types, tumors with the highest levels of both pathways’ transcripts were associated with longer survival (Fig. 3A-F). We previously noted an inverse correlation between mitochondrial β-FAO and glycolysis in HBs and HCCs and have attributed this to the preservation of the Randle cycle, whereby these two competing metabolic pathways engage in a negative feedback loop in response to metabolites such as ATP, AcCoA, malonylCoA and citrate as well as Cpt1a, phosphofructokinase and the pyruvate dehydrogenase complex (2,18,20,21,35,36). In two of the above cancer types, (lung adenocarcinoma and cervical squamous cell carcinoma) tumors expressing the lowest β-FAO- and ω-/peroxisomal-FAO-related transcript levels tended as a group to express higher levels of glycolysis-related transcripts (Fig. 3G). In contrast, clear cell kidney cancers associated with shorter overall survival expressed lover levels of glycolysis-related transcripts.

**Figure 3.**
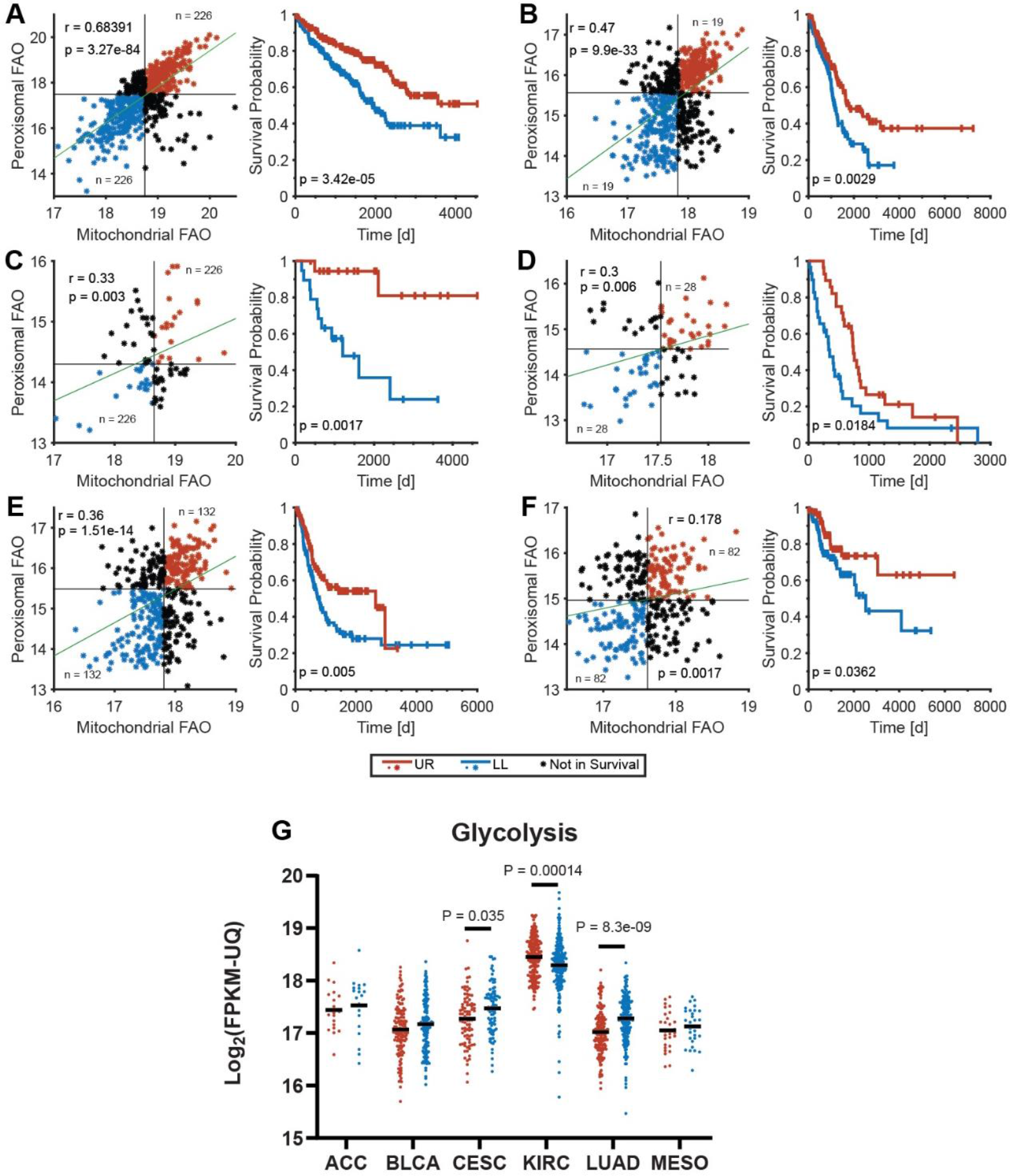
Mitochondrial β-FAO- and ω-/peroxisomal FAO-associated transcripts correlate directly with long-term survival in select cancers. Transcripts used were those from Fig. 2A&B and refs. (72,73). FPKM-UQ values were obtained from the GDC-PANCAN dataset via the UCSC Xenabrowser (xena.ucsc.edu). Average expression for each set of transcripts was determined from all 34 cancers in TCGA and expressed as a dot plot which was then divided into four quadrants, each containing approximately equal numbers of tumors. Long-term survival information was then obtained for each quartile of patients whose tumors’ transcript levels localized to the upper right and lower left portions of the dot plots. *A.* Kidney clear cell carcinoma, *B.* Lung adeno-carcinoma, *C*. Adrenocortical carcinoma, *D.* Mesothelioma, *E.* Bladder urothelial carcinoma. *F.* Cervical squamous cell carcinoma/endocervical adenoma carcinoma. *G.* The average expression of transcripts encoding glycolytic enzymes (72,73) was obtained from each of the above favorable and unfavorable survival cohorts. P values were determined using Welch’s t-test.

### The response to DDDA is associated with the induction of inflammatory markers and the development of DDDA resistance

The rapid onset of acute hepatic failure in *ehhadh-/-* mice provided DDDA-containing diets is associated with the induction of select transcripts encoding inflammatory markers and enzymes that participate in the Phase II anti-oxidant response and sphingomyelin biosynthesis (25). We evaluated several of these in the livers and HBs of mice maintained on standard or DDDA diets for 3 wks. In HBs from the DDDA diet cohort we noted significant increases in transcripts encoding macrophage migration inhibitory factor (Mif), the Phase II enzyme NAD[P]H:quinone oxidoreductase 1 (Nqo1) and the sphingolipid pathway enzymes delta 4-desaturase Degs1 and serine palmitoyl-transferase 2 (Sptlc2) (Fig. 4A). Together with the results presented in Fig. 1B and D, these findings indicate that, in response to DDDA, HBs up-regulate markers reminiscent of those detected in the livers of C12 diet-maintained *ehhadh-/-* mice (25).

**Figure 4.**
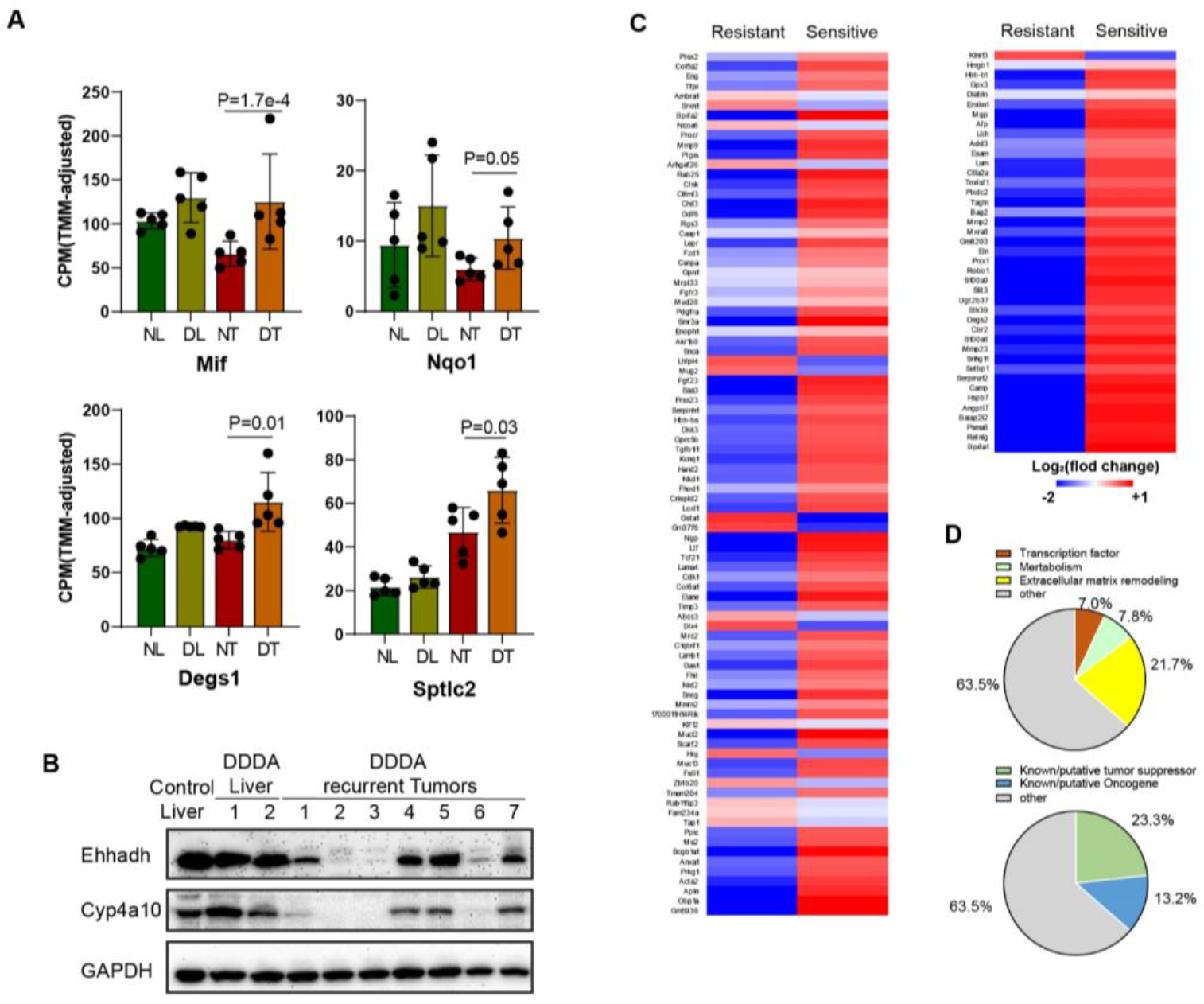
Inflammatory mediators and properties of DDDA-diet-resistant tumors. *A.* Levels of transcripts encoding the indicated inflammatory markers, Phase II anti-oxidants and sphingomyelin biosynthetic enzymes. *B.* Ehhadh and Cyp4a10 protein expression in livers and tumors from mice maintained on standard diets and similar to those shown in Fig. 2*C* were compared to those from mice maintained on DDDA diets for >20 wks. Four sets of tissues from each group were analyzed. *C.* RNAseq analysis on tumor groups maintained for three wks (Fig. 1A) or >20 wks on DDDA-supplemented diets. A total of 129 differences are shown here and includes both q<0.05 and q>0.05 but P<0.001 groups. See Supplementary Tables 2 and 3 for the identification and relative expression differences of these gene sets. *D.* Pie chart of the functional categories of transcripts depicted in *C* and Supplementary Tables 2 and 3.

Despite objective responses to both long- and short-term C12 and/or DDDA diets (Fig. 1), HB-bearing mice eventually succumbed to progressive disease (Fig. 1A and data not shown) with tumors no longer showing evidence of the widespread necrosis that characterized the initial response (Fig. 1B and C). Both Ehhadh and Cyp4a10 re-expression as a potential explanation for this acquired resistance were observed in some but not all tumors from mice maintained on DDDA diets for >20 wks (Fig. 4B). We therefore performed another RNAseq analysis on the resistant cohort that maintained low levels of Ehhadh and Cyp4a10 expression and compared the results to HBs that initially responded to a three week DDDA diet (sensitive cohort) with the intention of revealing the strategies employed by the former group to achieve DDDA resistance. To ensure inclusion of the broadest population of differentially expressed transcripts, we included those identified using any one of the three previously employed methods to quantify differential read counts, i.e. EdgeR, CLC Genomics Workbench and DESeq2 (q<0.05) and identified 41 differences. We then broadened the survey to include transcripts with q values >0.05 but P values <0.001 and identified an additional 88 differences (Fig. 4C and Supplementary Tables 1 and 2).

112 of the above 129 transcripts (86.8%) were down-regulated in the resistant cohort, with 23 of these (20.5%) being reduced >10-fold and only 9 (8.0%) being reduced <2-fold. In contrast, none of the 17 up-regulated transcripts was increased by >10-fold and 8 (47.0%) were increased <2-fold. Thus, the resistant HB cohort was characterized by disproportionate and more robust transcript repression. 28 (21.7%) of differentially expressed transcripts encoded structural components or remodelers of the extracellular matrix (ECM) or molecules functioning in cellular adhesion (Fig. 4D). Other prominent groups encoded transcription factors/co-activators (9 transcripts=7.0%) and enzymes regulating metabolism or redox balance (10 transcripts=7.8%). Perhaps most strikingly however, 32 (24.8%) of the transcripts encoded known or putative tumor suppressors, all but one of which were down-regulated in resistant tumors. An additional 17 (13.2%) of transcripts encoded known or putative oncoproteins or those which facilitate metastasis although only three of these were up-regulated. Thus, in total, >70% of the transcript differences between DDDA-sensitive and DDDA-resistant cohort encode proteins that likely alter tumor behavior, metabolism and extracellular environment.

The transcripts listed in Supplementary Tables 1 and 2 and their relationship to long-term survival were compiled from nearly 8000 samples of 17 different adult human cancers archived in The Human Protein Atlas (version 19.3) (https://www.proteinatlas.org/humanproteome/pathology). This showed that the levels of 63.6% of the transcripts (82 of 129) correlated with survival in one or more tumor types (Supplementary Table 3). In 145 of the 181 matches identified (80.1%), the correlation with survival was concordant, whereas in the remaining matches, it was discordant (P=1.78 x 10^−7^, Pearson’s Chi^2^ test). Additionally, the tumor types in which these correlations were observed were not random. For example renal cancers (KIRC, KIRP, KICH) and HCC (LIHC) were found to have the most frequent associations (92 and 22 examples, respectively) whereas only a single association or none at all was seen for gliomas, melanomas and cancers of the prostate and testes. These results indicate that DDDA-resistant HBs tended to down-regulate transcripts associated with long-term survival and up-regulate those associated with shorter survival.

### Effects of DDDA diets on liver and tumor metabolism

The normal liver relies more on β-FAO than glycolysis as its primary energy source whereas the reverse is true for HBs and HCCs (18,19,22,36). This is consistent with Warburg-type respiration but is not irreversible. For example, we have shown that this aberrant tumor metabolism can be at least partially normalized by high-fat diets (HFD), which restore liver-like levels of β-FAO, slow tumor growth and extend survival (36). This “forced metabolic normalization” likely reflects the preservation of Randle cycle regulation discussed above and the need for high rates of glycolysis to sustain tumor growth at its highest levels (2,35,36).

To determine whether DDDA diets might be contributing to HB inhibition via a similar mechanism, we compared oxygen consumption rates (OCRs) of partially purified mitochondria from the livers and tumors of the cohorts shown in Fig. 1A in response to several standard and anaplerotic TCA cycle substrates (Fig. 5A-C) (18,19,22,36). Consistent with our previous findings, total OCRs were significantly reduced in control diet tumors relative to control diet livers (Fig. 5A) with a somewhat greater contribution being made by Complex II in each case (Fig. 5B&C). This was consistent with the marked reduction in mitochondrial mass documented in murine HBs and HCCs (18,19,22,36). Unlike what had been previously observed for HFDs, which were comprised primarily of TCA cycle-compatible lipids such as palmitate (36), neither C12 nor DDDA exerted a discernible effect. Interestingly, however, each of two “outlier points” with the highest OCRs, originated from mice with the longest survival. A determination of mitochondrial mass in these two tumors, based on quantification of their mitochondrial DNA content, did not show any significant differences relative to other members of their cohort indicating that the normalization of their responses was due to a more efficient utilization of substrates.

**Figure 5.**
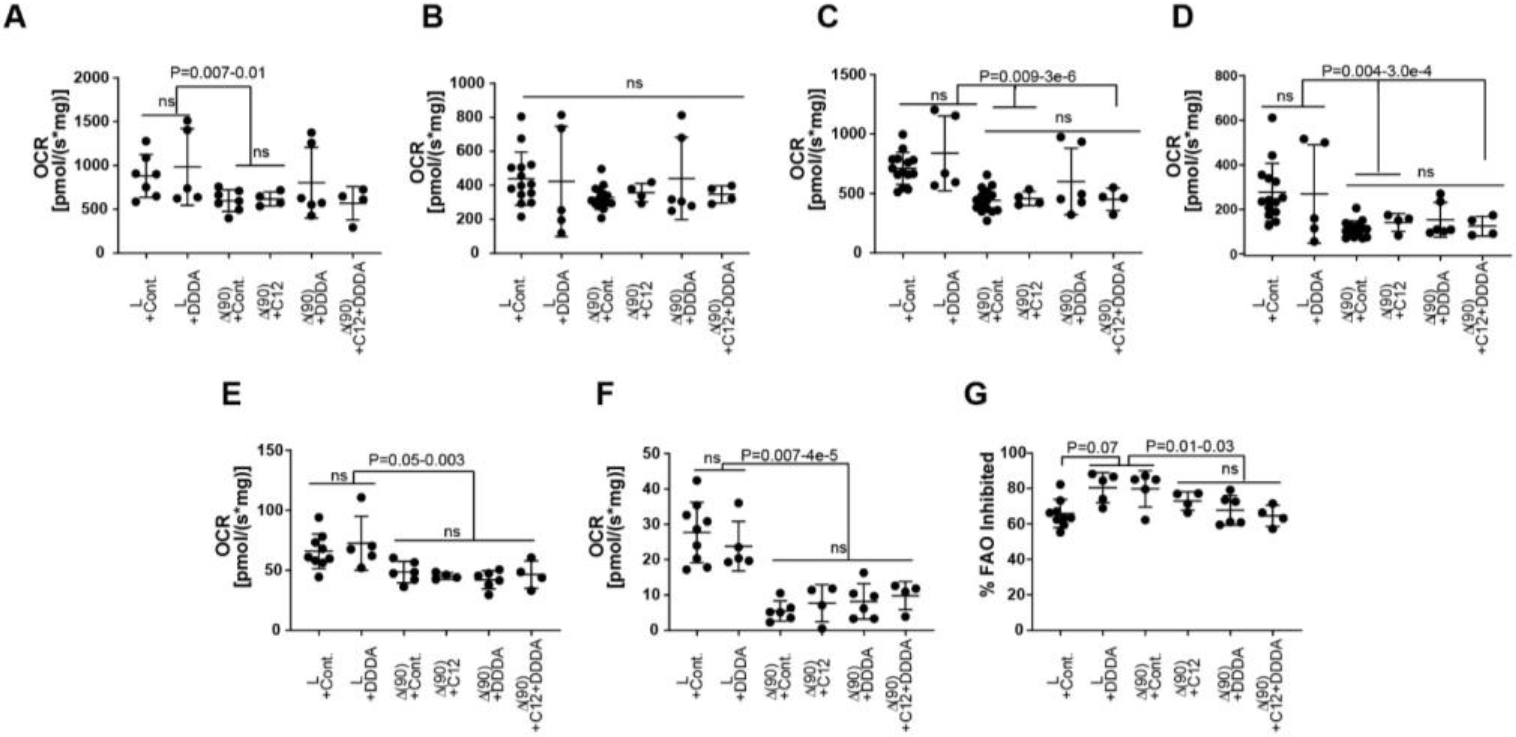
OCRs in livers and tumors from mice maintained on standard or DDDA-enriched diets. Mitochondria were isolated from livers and HBs of mice maintained on the indicated diets (Figure 2*A*) and OCRs were determined as previously described (18–22). *A.* Total Oxphos in response to the addition of cytochrome c, malate, ADP, pyruvate, glutamate and succinate. *B&C*. Complex I and Complex II activities activity. *D.* The glutamate response was determined after the addition of cytochrome c, malate, ADP and pyruvate. *E.* OCRs in response to palmitoyl-CoA were determined following the addition of cytochrome c, malate, ADP and L-carnitine. *F.* OCRs in response to L-carnitine was determined following the addition of cytochrome c, malate and ADP. *G.* The fraction of β-FAO that was Cpt1a-dependent was determined following the addition of etomoxir to palmitoyl-CoA- and L-carnitine-supplemented reactions as described in *E.*

Rather than increasing glutaminolysis as occurs in many tumors and cancer cell lines (1,5), HBs actually decrease this activity indicating that they are not reliant on the anaplerotic delivery of glutamate-derived α-ketoglutarate (19,22,36). This remained the case in HBs from mice on C12, DDDA and C12+DDDA diets (Fig. 5D).

Finally, we examined β-FAO in two ways. First, we measured the response to palmitoyl-CoA following the addition of all necessary substrates including L-carnitine (Fig. 5E). This allowed us to ascertain the inherent activity of mitochondrial β-FAO in a non substrate-limiting manner. Second, we measured the response to L-carnitine in the absence of exogenous palmitoyl-CoA as an indirect way of quantifying the pre-existing stores of fatty acyl-CoA derivatives that were available for immediate Cpt1a-mediated transport into the mitochondrial matrix (Fig. 5F). All tumors, regardless of the diets upon which the animals had been maintained, showed similar responses. Finally, the inhibition of Cpt1a by etomoxir showed that both livers and HBs, again irrespective of diet, were equally dependent upon active transport of medium-long chain fatty acids (Fig. 5G). In sum, these results strongly suggest that, despite the large-scale oxidation of C12 and DDDA by the ω-/peroxisomal-FAO pathway and its provision of TCA-substrates such as succinate (Fig. 2A), overall mitochondrial function in response to standard substrates was largely unaffected.

## Discussion

The optimization of tumor growth and survival in response to metabolic re-programming is well-documented and theoretically affords unique therapeutic opportunities, including those based on dietary manipulation (1,2,4,6,7,37–39).The latter include caloric restriction or glucose deprivation, which reduce Warburg-type respiration and temper insulin and/or insulin-like growth factor signaling (4,40–43). The ensuing changes in the micro-environmental nutrient supply can impact tumor cell signaling pathways, growth and chemotherapeutic sensitivity (9,44–46).

We previously exploited a mouse model of HCC in which the preservation of the Randle cycle allowed the suppression of glycolysis by β-FAO in response to HFDs, thereby impairing tumor growth and extending survival (36). A confirmatory analysis of primary human cancers showed that at least six tumor types with high β-FAO:glycolysis transcript ratios were also associated with significantly longer survival (36). Non-dietary approaches to achieving similar results include the direct pharmacologic inhibition of glycolysis, glutaminolysis and fatty acid synthesis or the specific targeting of mutant enzymes which generate onco-metabolites (14,15,47,48).

The common feature of the above approaches is that they restrict the production of metabolites that are produced at higher levels by transformed cells and thus serve to maximize tumor cell proliferation and/or block differentiation. For example, Warburg respiration produces glucose-6-phosphate, the initial substrate for the pentose phosphate pathway, as well as reducing equivalents in the form of NADPH. Downstream glycolytic reactions, in addition to providing ATP, generate 3-phosphoglycerate for serine, glycine and purine nucleotide biosynthesis and pyruvate, which links glycolysis and the TCA cycle but also furnishes alanine and the anaplerotic TCA cycle substrate oxaloacetate. Similarly, glutaminolysis anaplerotically provides α-ketoglutarate, another TCA cycle intermediate, that in turn supplies aspartate, asparagine and the citrate needed for *de novo* lipid biosynthesis (2,47,49). Finally the neomorphic activity of isocitrate dehydrogenase (IDH) missense mutants generates the novel oncometabolite 2-hydroxyglutarate, which maintains the undifferentiated state of the blast cell population in acute myelogenous leukemia (48,50). Inhibition of this mutant enzyme can induce terminal differentiation and hematologic remission (51,52).

A limitation of the above approaches is that they typically reduce tumor cell proliferation without necessarily eliminating the malignant clone. Compensatory metabolic adaptation can also circumvent such therapies by reducing the initial dependence on the targeted pathway (53). Cancer stem cells may also be less susceptible to such interventions either because of their inherent quiescence and/or because of different metabolic dependencies (16). Finally, pharmacologic targeting of cancer cell metabolic pathways is often limited by a narrow therapeutic window since the pathways being targeted are seldom unique and are invariably active in normal cells as well.

The approach taken in the current work differs from those described above in that it capitalizes on the tumor-specific loss of expression of a specific enzyme. This approach was inspired by the metabolic disorder Type I hereditary tyrosinemia, in which the absence of fumarylacetoacetate hydrolase (FAH) leads to the accumulation of the hepatotoxic metabolite fumarylacetoacetate (FAA) and ultimately to liver failure (54). Ehhadh deficiency is conceptually similar except that it is an acquired trait and thus strictly confined to the transformed hepatocyte population. The fact that both normal and tumor-bearing mice tolerate high dietary burdens of C12 and DDDA, either alone or in combination and without evidence of overt toxicity (25), speaks to a high therapeutic index that is difficult to achieve with other approaches.

Tumor inhibition by both C12 and DDDA (Fig. 1A) supports the idea that the toxicity is due to the accumulation of ω-/peroxisomal-FAO pathway-related metabolites-most likely DDDA itself-rather than to off-target effects. The fact that C12 and DDDA were not additive in extending survival also supports this idea and further indicates that the maximal anti-tumor effect is achieved with either compound. This was somewhat surprising in the case of C12 given the marked down-regulation of the two cytochrome P450 enzymes (Cyp4a10 and Cyp4a14) that participate in C12’s proximal catabolism (Fig. 2A-C). Sufficiently toxic levels of DDDA must therefore still accumulate in the face of C12-enriched diets despite the dramatic suppression of Cyp4a10 and Cyp4a14. The ensuing hepatotoxicity thus likely only requires levels of DDDA that can be generated by the reduced levels of these two cytochromes. The inability of C12 and DDDA to completely eliminate all tumor cells likely reflects their residual Ehhadh levels and/or their adaptation to otherwise toxic levels of DDDA (Fig. 2C&G).

The down-regulation of Ehhadh and other ω-/peroxisomal-FAO pathway-related transcripts seen in murine HBs (Fig. 2B) was recapitulated in Myc-driven murine HCCs, in human HBs and in subsets of several other human tumor types (Fig. 2D-F) and closely correlated with protein levels (Fig. 2C&G). In some cases, Ehhadh transcript down-regulation was particularly dramatic with some tumors showing >95% suppression relative to corresponding normal tissue (Fig. 2E). It is currently not clear whether this loss of Ehhadh in such a broad range of tumors would necessarily render them susceptible to C12 or DDDA or whether their toxic effects are hepatocyte-specific. In mice with germ-line *ehhadh* gene knockout, the most striking effect of dietary supplementation is massive hepatic necrosis with no toxicity having been reported in other tissues (25). However, it is certainly conceivable that the rapidity with which lethal hepatic failure develops could mask the protracted effects of C12 and DDDA elsewhere. Testing these compounds directly in non-hepatic cancers with documented down-regulation of the ω-/peroxisomal-FAO pathway will be needed to establish the broader efficacy of this dietary approach.

The direct relationship between mitochondrial β–FAO-and ω-/peroxisomal-FAO-related transcript down-regulation observed in murine and human HBs (Fig. 2B&D) was seen in other cancer types although, not unexpectedly, the correlations were somewhat more modest when tumors were considered as single homogeneous groups (Fig. 3A-F). However, in six cases, tumor subsets with the lowest levels of these transcripts were associated with shorter survivals relative to those with the highest levels. In three cancer types, an inverse relationship was seen between the transcripts and those encoding enzymes of the glycolytic pathway (Fig. 3G) but in only two of these were higher levels of glycolytic transcript expression associated with shorter survival. As already discussed with regard to HB, this inverse correlation between FAO and glycolysis is consistent with the preservation of Randle cycle-type regulation and agrees with previous reports demonstrating that survival for some cancers is inversely correlated with glycolytic pathway activity (55–61). The prolongation of survival by normalizing β-FAO with high fat diets and by reciprocally suppressing Warburg-type respiration (36) suggests that this approach could potentially be combined with C12 or DDDA-enriched diets to capitalize on the additional ω-/peroxisomal-FAO pathway vulnerability described here.

The eventual progression of tumors in all DDDA-treated mice, despite impressive initial responses (Fig. 2B-D) was in keeping with numerous other studies involving dietary interventions administered either singly or in combination with chemotherapeutic agents (4,36,40,42,62,63). Resistant HBs that failed to re-express Ehhadh and Cyp4a10 (Fig. 4B) altered the expression of only 129 genes, nearly 90% of which were down-regulated relative to those of DDDA-sensitive tumors (Fig. 4E and Supplementary Tables 1 and 2). Despite its small size, this collection may be an over-estimate since we used three independent genomic analysis methods to identify its members and did not require that they necessarily agree. In some cases, we also liberalized the FDR and instead relied exclusively on P values of <0.001. Given these caveats, perhaps the most surprising finding of this analysis was the nature of the gene expression differences between the two tumor groups. Only 10 transcripts encoded metabolic enzymes or those functioning to alleviate oxidative/electrophillic stress or redox imbalances. This suggested that the development of this form of DDDA resistance might involve post-transcriptional changes in the activities and/or levels of enzymes lying outside the ω-/peroxisomal-FAO pathway depicted in Fig. 2A and that might serve to more efficiently reduce intracellular DDDA levels. This might involve changes in the ECM as evidenced by the altered expression of 28 transcripts encoding several of its components, including enzymes such as matrix metallo-proteases or several cell-cell or cell-matrix adhesion molecules. Such changes often affect chemotherapeutic sensitivity and could impact the efficiency with which DDDA traverses the vasculature and enters tumor cells (64–69). The down-regulation of transcripts encoding endothelial cell-related proteins such as Eng, Angptl7, Esam and Scarf2 might further contribute to alterations in blood vessel permeability.

\More than half the transcripts listed in Supplementary Tables 1 and 2 correlated with long-term, favorable survival among a group of nearly 8000 samples representing 17 general cancer types from TCGA (Supplementary Table 3). These associations were concordant in nearly 80% of the cases and were particularly prominent in renal cancers and HCCs. Combined with the selective differential expression of transcripts functioning in transcriptional regulation, metabolism and extracellular environment, these findings strongly suggest that resistant tumors represent a more aggressive form of HB that arises as a consequence of the reprogramming of a limited and functionally diverse set of pathways.

In most cases, C12 and/or DDDA-supplemented diets did not affect the OCRs of livers or tumors in response to different substrates (Fig. 5). This is perhaps to be expected given that peroxisomal β-FAO does not generate energy directly although it does provide select TCA cycle substrates such as succinate and AcCoA (Fig. 2A) (31). Two notable outliers to this, however, were seen in response to the full complement of non-fatty acid-related TCA cycle substrates (Fig. 5A-C) and both of these were associated with mice having the longest survival. Mitochondria from these two tumors showed OCR responses at the upper end of the range seen in control livers and were reminiscent of those seen in livers or in tumors from animals maintained on high fat diets (36). However, the mitochondrial DNA content of these tumors was not increased relative to that of the other tumors. The basis for this apparent normalization and the degree to which it affects survival is currently unknown although it provides additional reason to believe that a modification of our previously used HFD regimen so as include C12 might provide additional survival benefit by targeting both mitochondria and peroxisomes (36).

Collectively, our results show that the severe acquired Ehhadh deficiency of HBs, HCCs and certain other cancers represents a distinct, tumor-specific and actionable metabolic susceptibility. The selective reduction of this enzyme causes the aberrant accumulation of DDDA, a highly toxic product of ω-/peroxisomal-FAO (25), and extensive tumor cell killing. At the same time, normal liver parenchyma and other tissues are spared from any discernible toxicity as they maintain levels of Ehhadh sufficient to metabolize DDDA (Fig. 2A). The ability to achieve significant anti-tumor responses with DDDA and/or C12 potentially offers simple, inexpensive, well-tolerated and non-cross-resistant therapeutic options that would be compatible with more traditional treatments and for individuals who were not candidates for or otherwise refused them.

## Experimental procedures

### HB generation and diets

HDTVIs of SB vectors encoding β-catenin and YAP^S127A^ were performed as previously described in 6-7 wk old FVB mice (Jackson Labs, Bar Harbor, ME) (19–21,70). Mice were monitored weekly and at least thrice weekly after tumors first became detectable. Animals were sacrificed when tumors reached a size of 2 cm in any dimension or when signs of stress were observed. All injections, monitoring procedures and other routine care and husbandry were approved by The University of Pittsburgh Department of Laboratory and Animal Resources and the Institutional Animal Care and Use Committee (IACUC). Diets consisted of standard animal chow which, where indicated, were supplemented with 10% (w/w) C12 in the form of trilauryl glycerate and/or 10% dodecandedioic acid (DDDA) (Sigma-Aldrich, Inc. St. Louis, MO). Tumor size and appearance were monitored by The UPMC Children’s Hospital Core Animal Imaging Facility using a 7-Teslar micro-MRI system (Bruker, BioSpin 70/30).

### Immunoblotting

Tissue lysates were prepared for immune-blotting as previously described (18,19). Antibodies used included those against Ehhad (1:500) (#ab93173, Abcam, Inc. Cambridge, MA), Cyp4a10 (1:1000) (#PA3-033, Invitrogen/Thermo-Fisher, Inc, Waltham, MA) and GAPDH (1:10,000) (Sigma-Aldrich, St. Louis, MO).

### Respirometry/metabolic studies

Oxygen consumption rates (OCRs) were performed on 50-100 mg of disrupted tissue immediately after sacrifice as previously described (18–21,36,71). Studies were performed with an Oroboros Oxygraph 2k instrument (Oroboros Instruments, Inc., Innsbruck, Austria) (18,19,71). Baseline OCRs were determined in 2 ml samples in Mir05 buffer following the addition of cytochrome c (final concentration=10 μM), malate (2 mM), ADP (5 mM), pyruvate (5 mM) and glutamate (10 mM). Together, these substrates provided and estimate of the activity of Complex I. Succinate was then added (10 mM final concentration) to determine the additional contribution provided by Complex II. The Complex I inhibitor rotenone was then added (0.5 μM final concentration) to calculate the proportional contributions of Complexes I and II. Activities were normalized to total protein.

### Quantification of transcripts encoding ω-/peroxisomal-FAO in human cancers

Expression values (FPKM-UQ) for all relevant transcripts were down-loaded from the GDC-PANCAN dataset via the UCSC Xenabrowser (xena.ucsc.edu). In cases where a patient had more than one sample of the same type, the expression values were averaged. Significance of expression differences between matched primary tumor and normal tissue samples was assessed for each cancer using Welch’s t-test where the members of the groups were the base-two logarithms of the expression values for each patient. The average tumor to liver expression ratio for each cancer was calculated by taking the ratio of the average expression across all tumor samples and dividing by the average for normal tissue samples.

## Data availability

Raw and processed RNA-Seq data were deposited in the National Center for Biotechnology Information (NCBI) Gene Expression and are accessible through GEO (https://www.ncbi.nlm.nim.gov/geo) at accession number GSE156545. All remaining data are contained within the article.

## Acknowledgements

We thank Jerry Vockley for valuable discussions and suggestions. This work was supported by NIH grants RO1CA174713 to EVP and DK090242 to ESG.

## Author contributions

Conceived experiments: ESG, HW, EVP

Performed experiments: HW, XC, MS, JG, JW, JL, SD

Analyzed and interpreted data: HW, JL, JAM, EVP

Wrote the manuscript: HW, EVP

## Supplementary Figure Legends

**Supplementary Figure 1.**
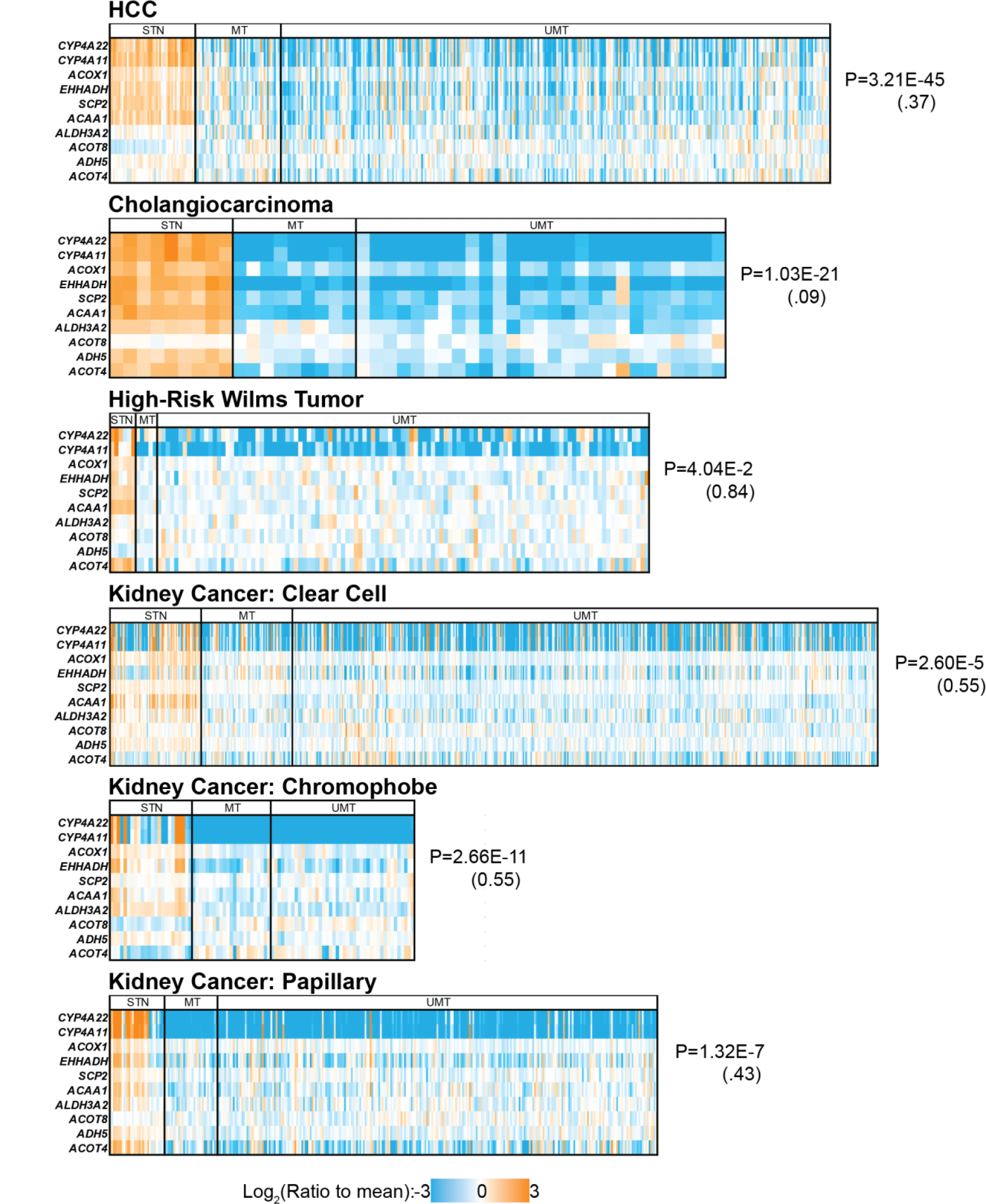
Down regulation of the ω-/peroxisomal-FAO pathways in human cancers. Expression of **ω-**/peroxisomal-FAO pathway transcripts are shown in primary tumors from the indicated cancers. Matched primary tissues are shown at the extreme left of each heatmap. Numbers to the right in parentheses indicate the average extent of down-regulation of all tumors and P values relative to the expression of the corresponding transcripts in matched control tissues.

## Supplementary Tables

**Supplementary Table 1.**
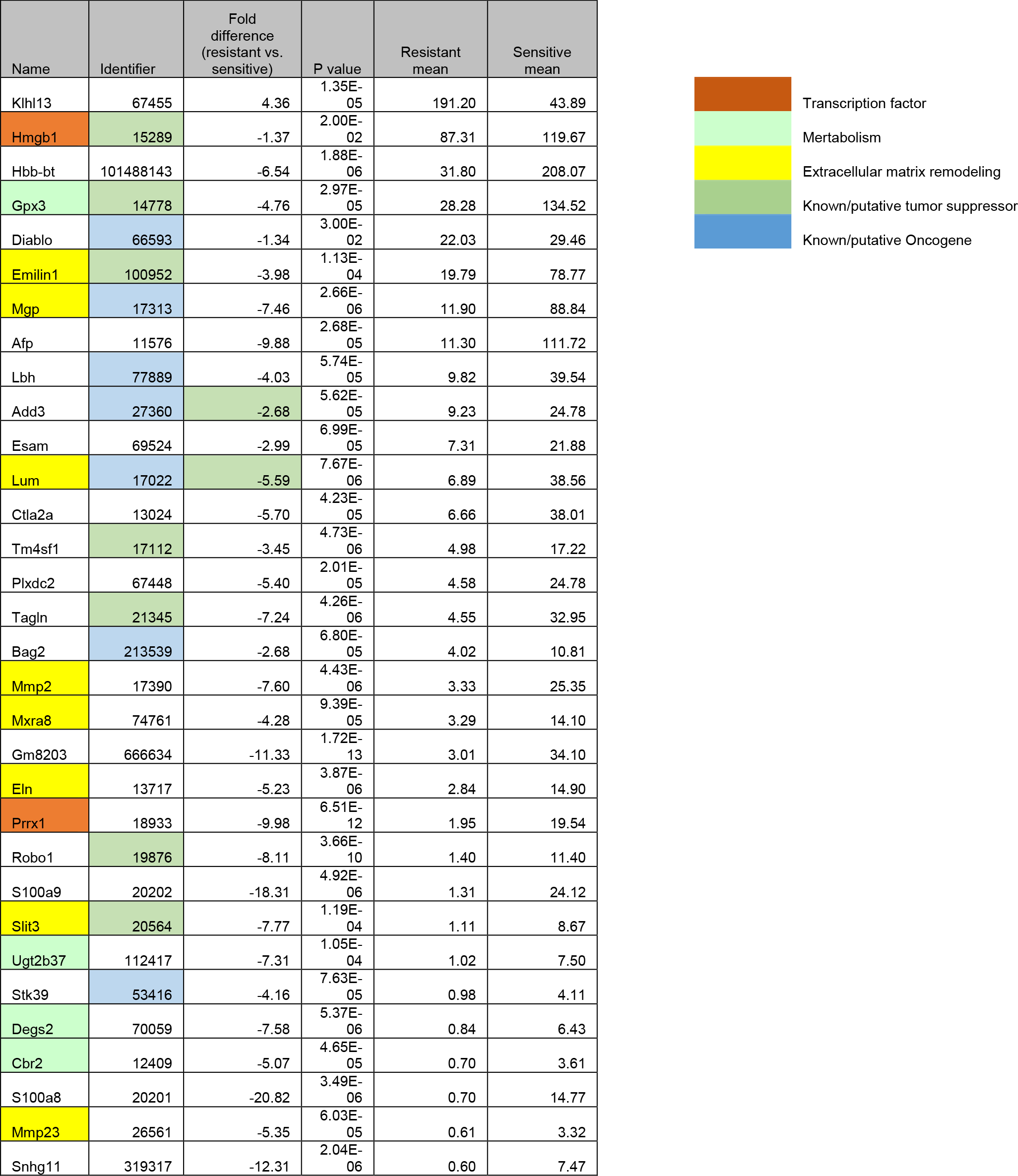
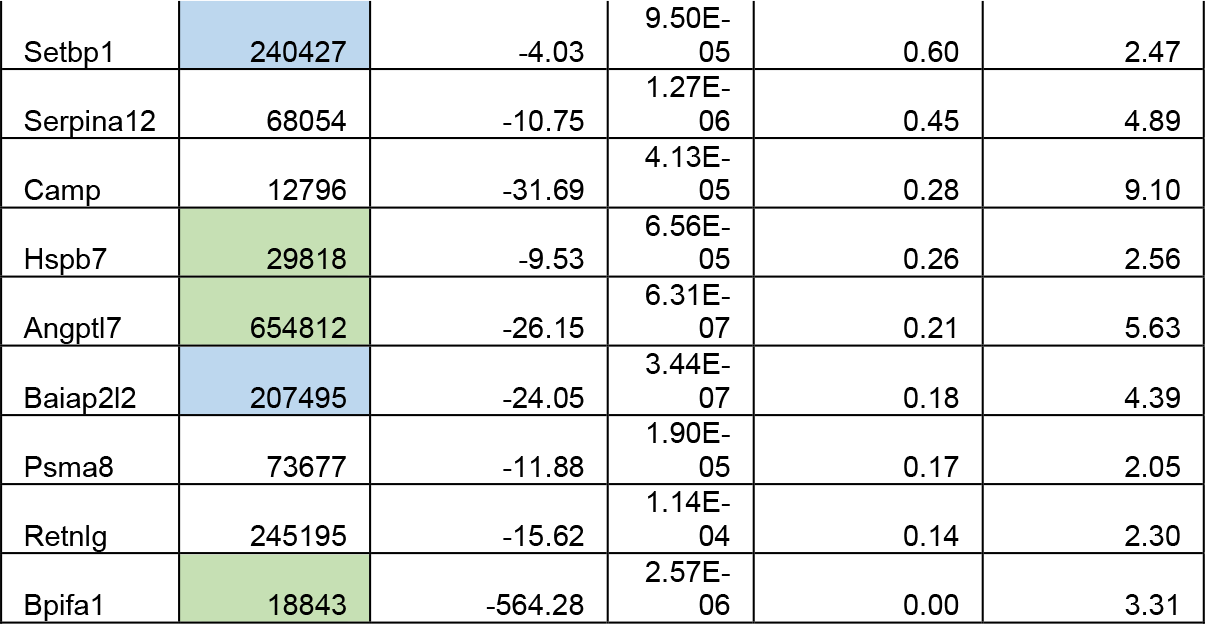
Gene expression differences in HBs arising in mice maintained on DDDA diets for 3 wks (sensitive group) versus 20-25 wks (resistant group). q<0.05 in all cases. Negative values indicate down-regulation in the latter group relative to the former group. In the first column, transcripts encoding proteins involved in extracellular matrix architecture and its re-modeling or cell adhesion are highlighted in yellow, those encoding transcription factors or co-activators are highlighted in red and those encoding enzymes that regulate metabolism or maintain redox homeostasis are highlighted in aqua. In the second column, transcripts encoding known or putative tumor suppressors are highlighted in blue and those encoding known or putative oncoproteins are highlighted in green. Note that dual functionality has been attributed to two transcripts (Add3 and Lum).

**Supplementary Table 2.**
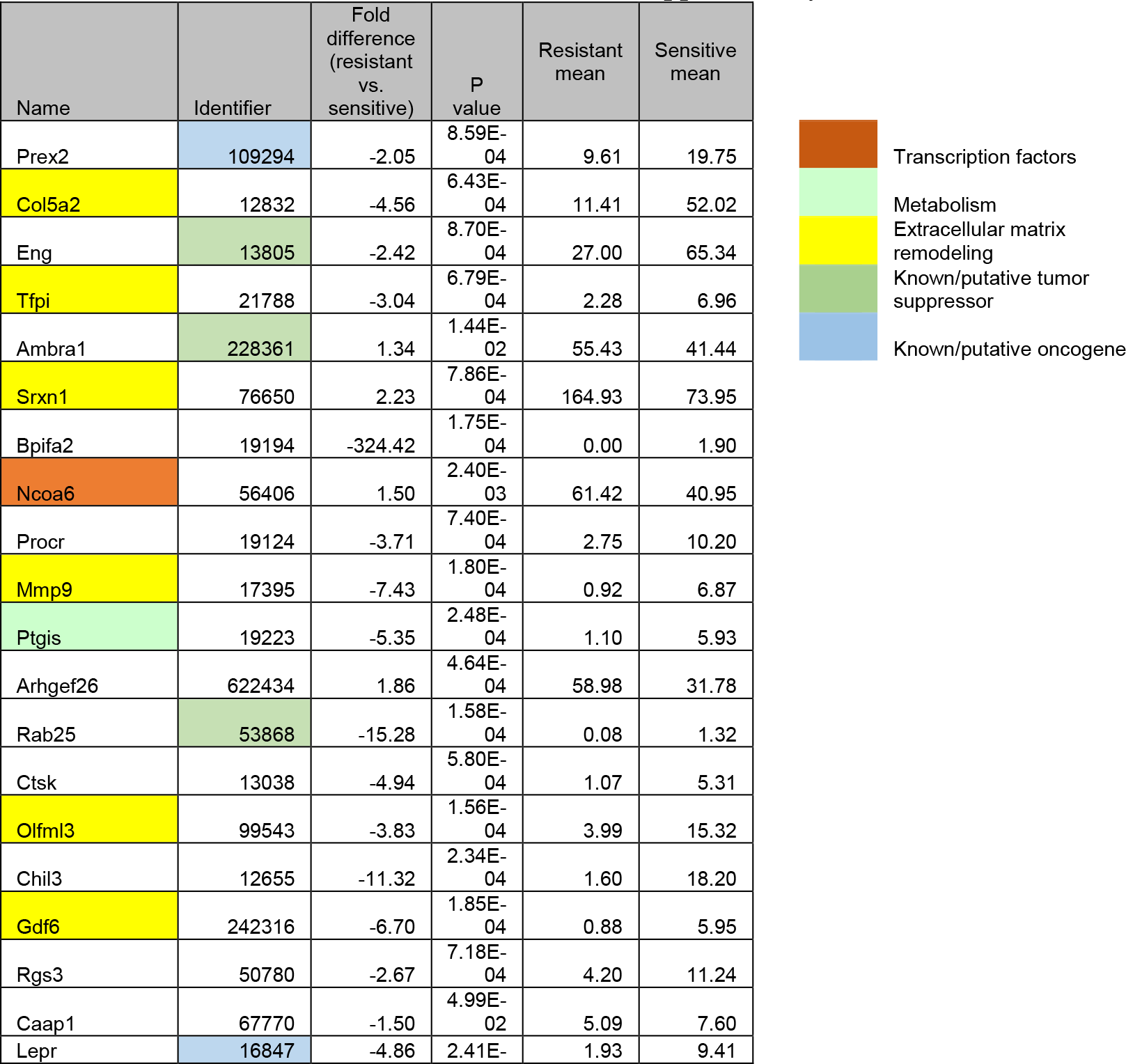
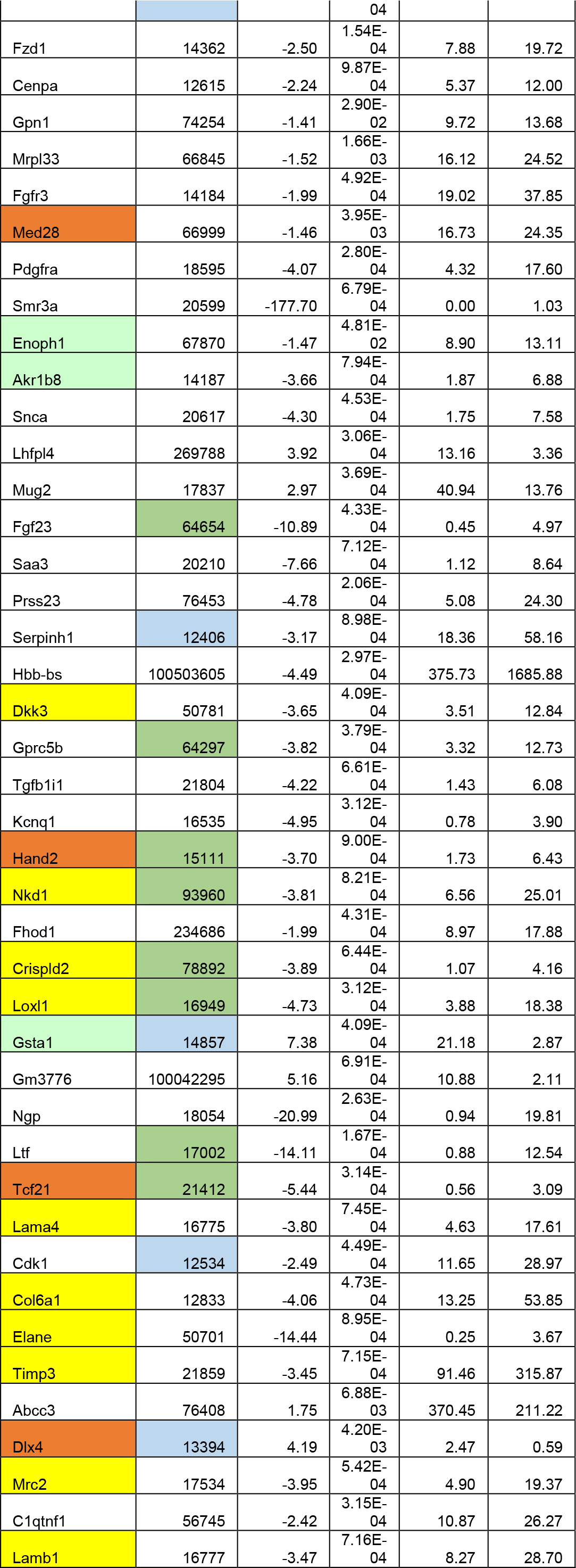
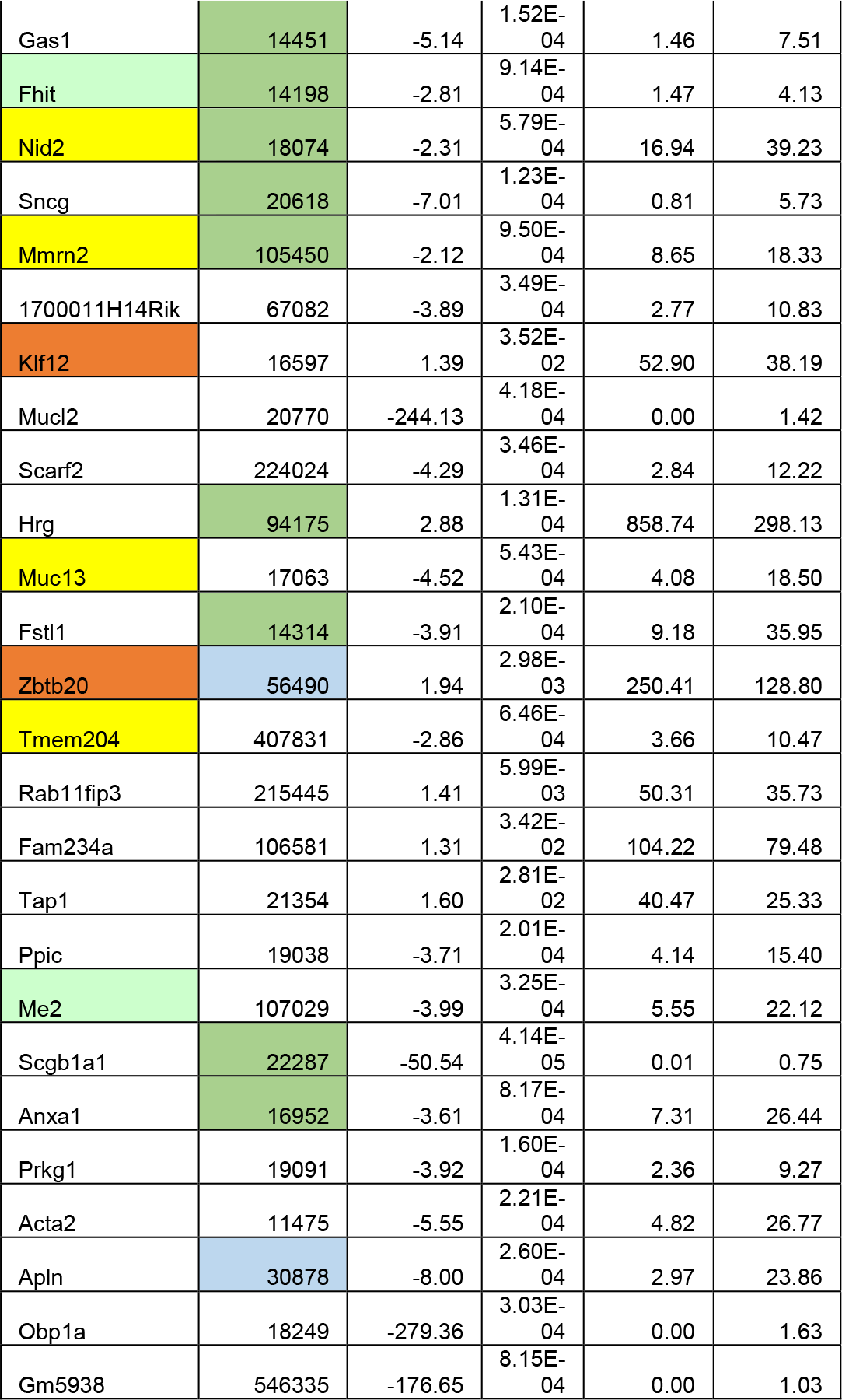
Additional gene expression differences in HBs arising in mice maintained on the diets described in Supplementary Table 2. In all cases, q>0.05, P<0.001. Transcripts encoding proteins involved in extracellular matrix architecture and maintenance or cell adhesion are highlighted in yellow, those encoding transcription factors or co-activators are highlighted in red and those encoding enzymes that regulate metabolism or maintenance redox homeostasis are highlighted in aqua. Highlighted color schemes are the same as those described in Supplementary Table 1.

**Supplementary Table. 3.**
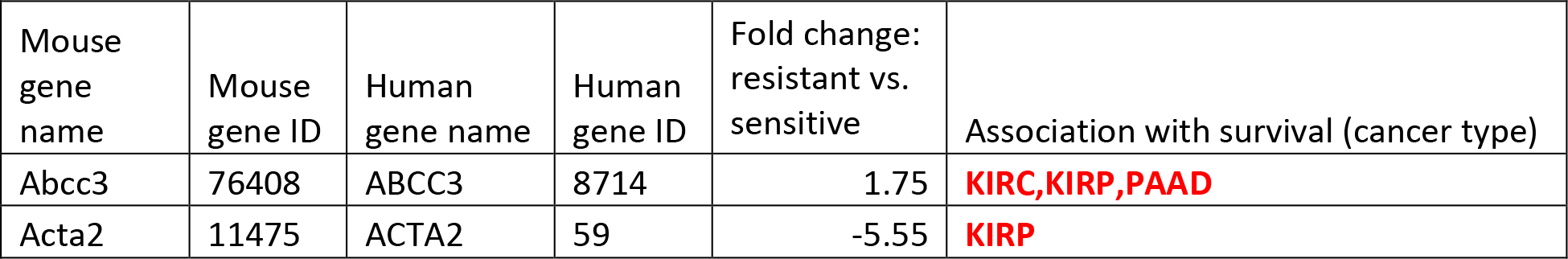
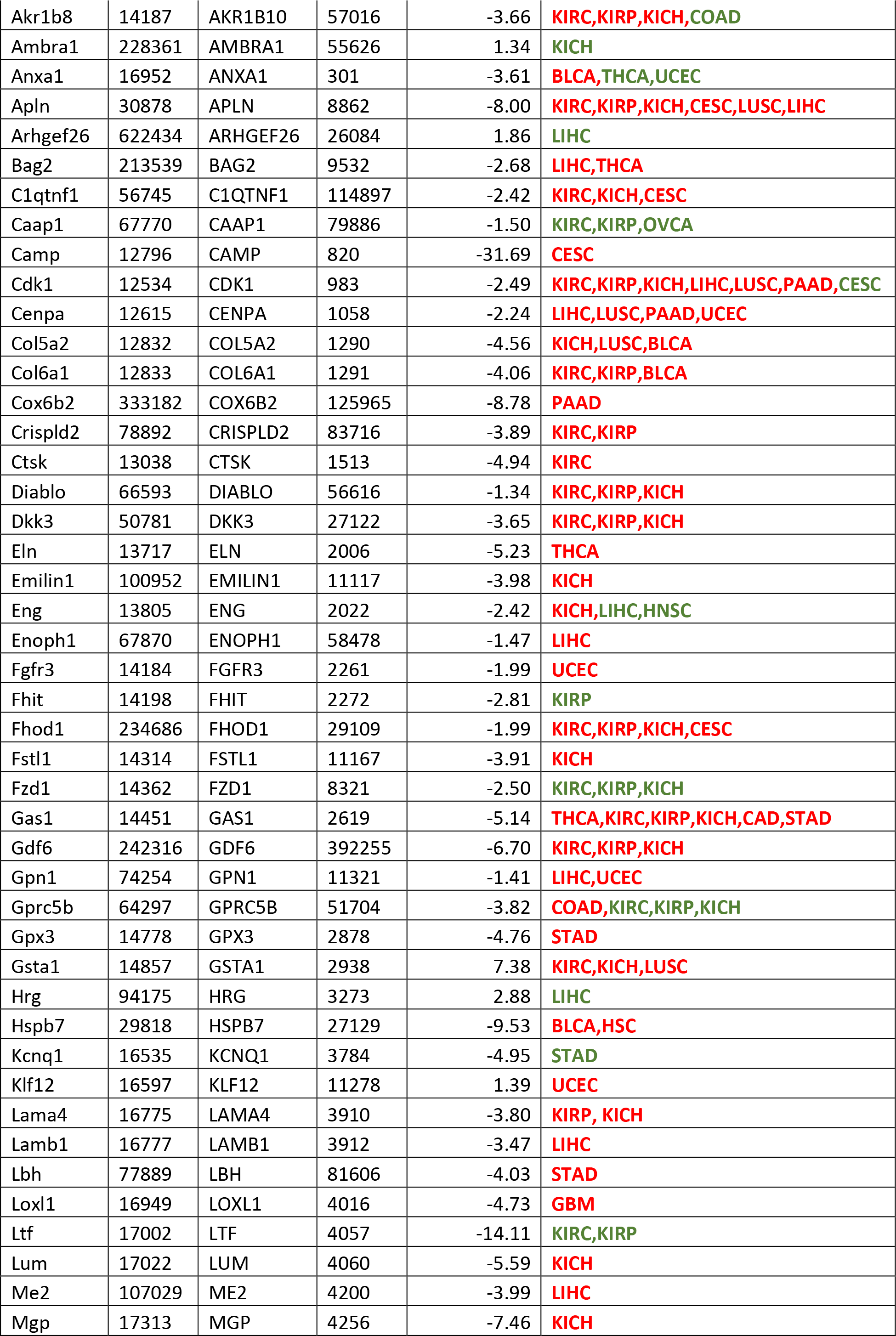
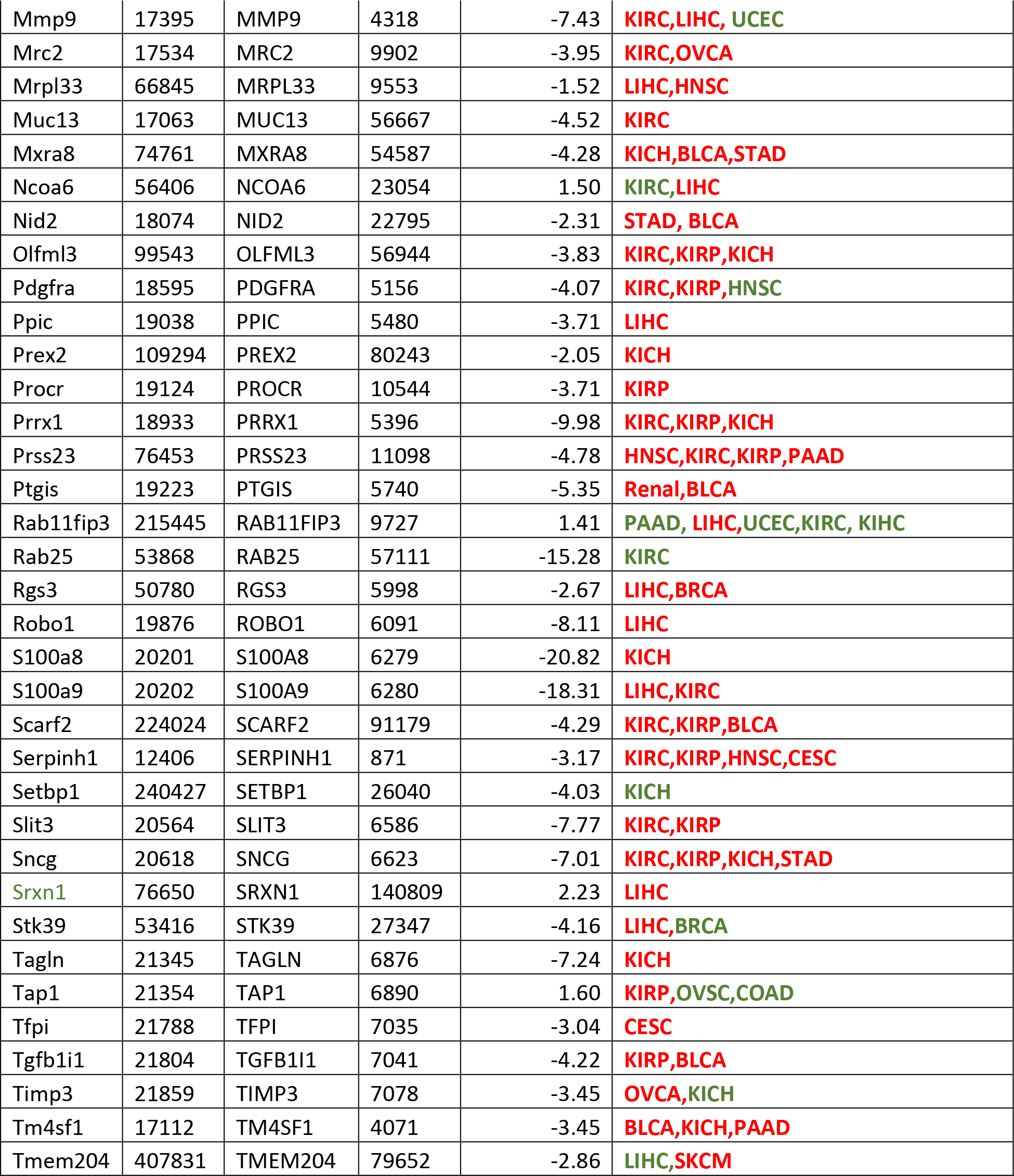
Correlation of gene expression differences from resistant tumors with survival in human cancers. Shown here are the genes from Supplementary Tables 2 and 3 whose expression correlated with survival in the cancer types listed in The Human Protein Atlas (https://www.proteinatlas.org/humanproteome/pathology). Tumors indicated in red are those in which low transcript expression correlated with significant shorter survival; those in green are those in which low transcript levels correlated with longer survival.

